# Spatial and temporal partitioning and tree preference in California woodland ants

**DOI:** 10.1101/745034

**Authors:** Dylan MacArthur-Waltz, Rebecca Nelson, Gail Lee, Deborah M. Gordon

## Abstract

Spatial and temporal partitioning of habitat may facilitate diversity and have important impacts on ant communities. To investigate niche overlap in an ant community in a northern California oak woodland, we observed ant foraging on trees in 4 seasonal surveys, each lasting 2 weeks, in a 9.5-hectare plot over the course of a year. Foraging activity in all 5 observed ant species differed by season, time of day, and/or the genera of trees used. Of the 3 ant species most frequently observed, *Camponotus semitestaceus* was most active during spring and summer nights, *Formica moki* was most active during spring and summer days, and *Prenolepis imparis* was most active during both day and night of fall and winter. All ant species preferred native trees to exotic trees and preferred evergreen trees to deciduous trees. Our results suggest that native evergreen oaks such as *Quercus agrifolia*, currently threatened by sudden oak death (*Phytophthora ramorum*), may be important for supporting ant biodiversity.

## Introduction

A central question in community ecology is how communities preserve diversity despite interspecific competition (1–3). Ant communities are ideal systems for studying coexistence because they are diverse, may include many similar species (which are theoretically more likely to compete strongly), and greatly influence ecosystem function (4–6). Many ant species have strong associations with particular host plant species (6–8), and this may reduce competition among coexisting ant species.

Competing ant species exhibit niche partitioning in foraging activity along many different axes, including season (9–11), day and night (4,9,10,12–14), temperature range (15,16), and space (12,17). With some exceptions (9,10), few studies have examined how seasonal and diel scales interact to influence ant community structure.

Urban and fragmented ecosystems may create complex, patchwork habitats that allow some groups of organisms to partition niches effectively (18). Native ant biodiversity in northern California is strongly linked to urbanization; disturbed, urbanized habitats have lower native ant species richness but support higher levels of invasive ant species. (19). Many species of ants rely on the honeydew produced by hemipteran insects as a nutritious food source (20–22). Compared to native trees, non-native trees have been shown to provide an inferior food source for ants because they harbor, on average, a lower diversity and abundance of hemipterans (23).

In our study system, five common native ant species (*Camponotus semitestaceus, Formica moki, Liometopum occidentale, Prenolepis imparis*, and *Tapinoma sessile*) coexist in a mixed native-exotic woodland in northern California in a Mediterranean climate with wet winters and dry summers. All five ant species forage on trees and all apparently forage on aphid honeydew (24–26).

Our study site has a mixture of native and exotic tree species planted over the last 200 years; previously the site was a mixture of grassland and oak savanna (27). Typical native tree species include oaks: *Quercus agrifolia*, *Q. lobata*, and *Q. douglasii*; typical exotic species include *Eucalyptus spp*., *Olea europaea*, and *Schinus molle*. Oaks play an important role in maintaining biodiversity in California forests (28). *Quercus agrifolia* is vulnerable to sudden oak death (*Phytophthora ramorum*), a pathogen of emerging concern in northern California (28). Disturbances such as sudden oak death may negatively affect ant community dynamics.

We conducted seasonal surveys during both day and night, across a variety of tree genera, to examine niche partitioning in an urbanized, northern Californian ant community. We asked the following questions:

1. Do ant species differ in foraging activity across seasons?
2. Do ant species differ in foraging activity between day and night?
3. Do ant species differ in their use of tree genera, native versus exotic trees, and deciduous versus evergreen trees for foraging?

## Methods

### Survey Region

We surveyed ant activity on and around tree trunks in a 9.53 ha region of the wooded Arboretum on the Stanford University campus. The survey region consisted of mixed oak-exotic woodland without significant brushy understory. Trees were identified using maps from the Stanford Maps and Records office (340 Bonair Siding Rd., Stanford, CA), which included GPS coordinates and species for each tree. Common ant species at the site included *Camponotus semitestaceus, Formica moki, Liometopum occidentale, Prenolepis imparis*, and *Tapinoma sessile*. We did not observe any other ant species at our study site in significant abundance (i.e., inhabiting >10 trees in any season). We observed *Linepithema humile* during spring, summer, and fall surveys in <10 trees on the northwestern edge of the survey region.

Four seasonal surveys, each lasting up to 2 weeks, were conducted in 2016-2017 from 29-July- 2016 – 8-August-2016 (Summer); 12-November-2016 – 13-November-2016 (Fall); 22-February-2017 - 26-February-2017 (Winter); and 12-May-2017 – 19-May-2017 (Spring). We surveyed each of approximately 870 trees twice during each season, once during the day and once at night, for a total of 7008 individual tree observations. We did not survey on rainy days as ants were not active. We did not survey in the hour before or after sunrise and sunset. We eliminated trees that died between surveys from the sample set. For about 20 trees, growth around the base was too dense to allow observation, and those trees were not included in our final analysis.

Observations were made by the authors and six research assistants. Ants were identified to species in the field or after observations using specimens collected at the time of observation. During each survey, we observed the bottom 2m of the tree trunk and a 1m radius around the base of the tree. We recorded all ant species present on the trunk and the base, and we estimated their abundance as follows: low abundance = 1-5, medium abundance = 6-30, high abundance =>30 individuals. We also recorded the presence of trails of any ant species. A trail was recorded if ants appeared concentrated in a line on the tree trunk (as opposed to scattered randomly around the trunk) or if ants moving in opposite directions touched antennae in passing. Observations lasted approximately 30 seconds - 1 minute per tree. All data are available for download (29).

We observed but did not map the presence of nests within the survey region. We observed about 10 nests of *Liometopum occidentale* only in *Quercus agrifolia*. We observed several *Prenolepis imparis* nests on the ground, greater than 10 *Camponotus semitestaceus* nests on the ground and 2 *Formica moki* nests on the ground. We did not find any *Tapinoma sessile* nests.

### Data Analysis

All statistical tests were performed in RStudio Version 1.1.383. We compared ant presence across seasons, day/night, tree genera, and ant species using Kruskal-Wallis rank sum tests. We then performed a second series of Kruskal-Wallis tests with the same factors and ant abundance as the response variable, using the median values of our bins of ant abundance in each category (1-5 ants ≅3, 6-29 ants ≅18, 30+ ants ≅100). Because the results of the presence/absence model and the abundance model were very similar (Table S1), we then used only the presence/absence model for species-specific analysis.

Season*Day_Night interaction terms were significant for the majority of species observed, so we combined Season (four levels) and Day_Night (two levels) into a single factor with eight levels (for example, one level was Summer Day). We ran post hoc Dunn tests on this combined variable for each ant species. We adjusted all p-values using the Benjamini-Hochberg procedure.

For each of the 5 most common ant species: *Camponotus semitestaceus, Formica moki, Liometopum occidentale, Prenolepis imparis*, and *Tapinoma sessile*, we performed a Kruskal-Walls tests on season, day/night, and genus individually. We used post hoc Dunn tests to determine the effect of season, day/night, and tree genus preference on ant presence for each individual species.

For each survey, we calculated the percentage of all deciduous, evergreen, native, and non-native trees that were occupied by any species of ant. We asked whether all species of ants preferentially used deciduous or evergreen trees, and native or exotic trees, using paired t-tests.

## Results

### Question 1: Do ant species differ in foraging activity across seasons?

The species present depended on season (Kruskal-Wallis, χ^2^=265.6, p < 2.2×10^-16^). *Camponotus semitestaceus* was observed most frequently during the spring and summer. *Formica moki* was observed most frequently during the spring and summer. *Liometopum occidentale* was observed most frequently during the summer. *Prenolepis imparis* was observed most frequently in the fall, slightly less frequently in the winter, less in the spring, and rarely in the summer. *Tapinoma sessile* was observed most frequently during the summer (See Figure 1 for Dunn post hoc results).

**Figure 1.**
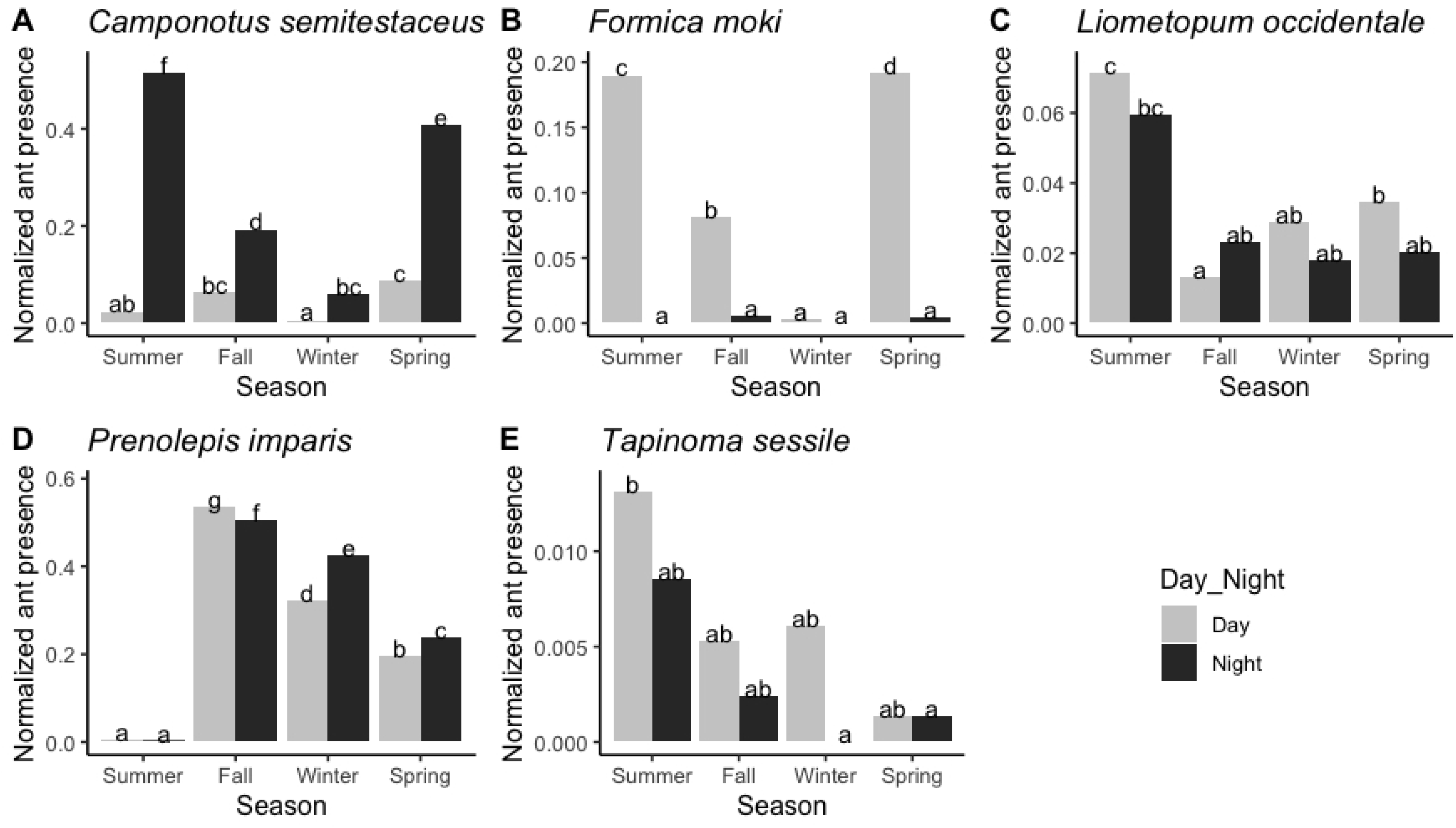
Normalized ant presence by season and day/night for: (a) *Camponotus semitestaceus*; (b) *Formica moki;* (c) *Liometopum occidentale;* (d) *Prenolepis imparis;* (e) *Tapinoma sessile*. Normalized ant presence is the number of trees on which an ant species was present divided by the total number of trees surveyed. Gray bars represent observations for day, and black bars represent observations for night. Letters show significant differences in Dunn tests at the p < 0.05 level.

### Question 2: Do ant species differ in foraging activity between day and night?

Most species differed in activity between day and night (Kruskal-Wallis, χ^2^=92.0, p < 2.2×10^-16^, Table S1), and these day-night differences varied among seasons. *Camponotus semitestaceus* was significantly more active at night than during the day, and this activity level depended on season (Kruskal-Wallis, χ^2^=1660, p < 2.2×10^-16^). *C. semitestaceus* was most often observed during spring and summer nights and observed less commonly during fall and winter nights (Figure 1A). During all seasons except winter, *Formica moki* was more active in the day than at night, and this activity level depended on season (Kruskal-Wallis, χ^2^=897.98, p < 2.2×10^-16^). *F. moki* was most often observed on spring and summer days (Figure 1B). Activity in *Liometopum occidentale* depended both on time of day and season (Kruskal-Wallis, χ^2^=59.646, p < 1.776×10^-10^). *L. occidentale* was more active on summer days than any time during fall, winter, and spring (Figure 1C). *Prenolepis imparis* was more active during the night in winter and spring and more active during the day in fall (Kruskal-Wallis, χ^2^=1483.9, p < 2.2x 10^-16^, Figure 1D). Activity in *T. sessile* depended on a time of day/season interaction (Kruskal-Wallis, χ^2^=19.0, p = 0.0083, Figure 1E). All post hoc Dunn test results are presented in Figure 1, and Kruskal-Wallis results are presented in Tables S1 and S2.

### Question 3: Do ant species differ in their use of tree genera, native versus exotic trees, and deciduous versus evergreen trees for foraging?

Species differed in their use of tree genera (Kruskal-Wallis, χ^2^=216.0, p < 2.2×10^-16^); this effect was observed in all species except *Tapinoma sessile* (Kruskal-Wallis test; CS: χ^2^= 66.3, p < 2.4×10^-12^; FM, χ^2^= 54.1, p < 7.0×10^-10^; LO: χ^2^= 63.1, p < 1.0×10^-11^; PI: χ^2^=182.8, p < 2.2×10^-16^; TS: χ^2^= 5.8, p = 0.44, Figure 3).

*Camponotus semitestaceus* and *Formica moki* were relatively uncommon on *Olea* trees (Figure 2A, 2B), while *P. imparis used Olea* trees more often than any other tree genus (Figure 2D). *C. semitestaceus* preferred *Quercus* and *Eucalyptus* trees over *Olea* and *Schinus* trees (Figure 2A). *F. moki* used most tree genera rather evenly but avoided *Olea* trees (Figure 2B). *Liometopum occidentale* preferred *Cedrus* trees and then *Quercus* (Figure 2C).

**Figure 2.**
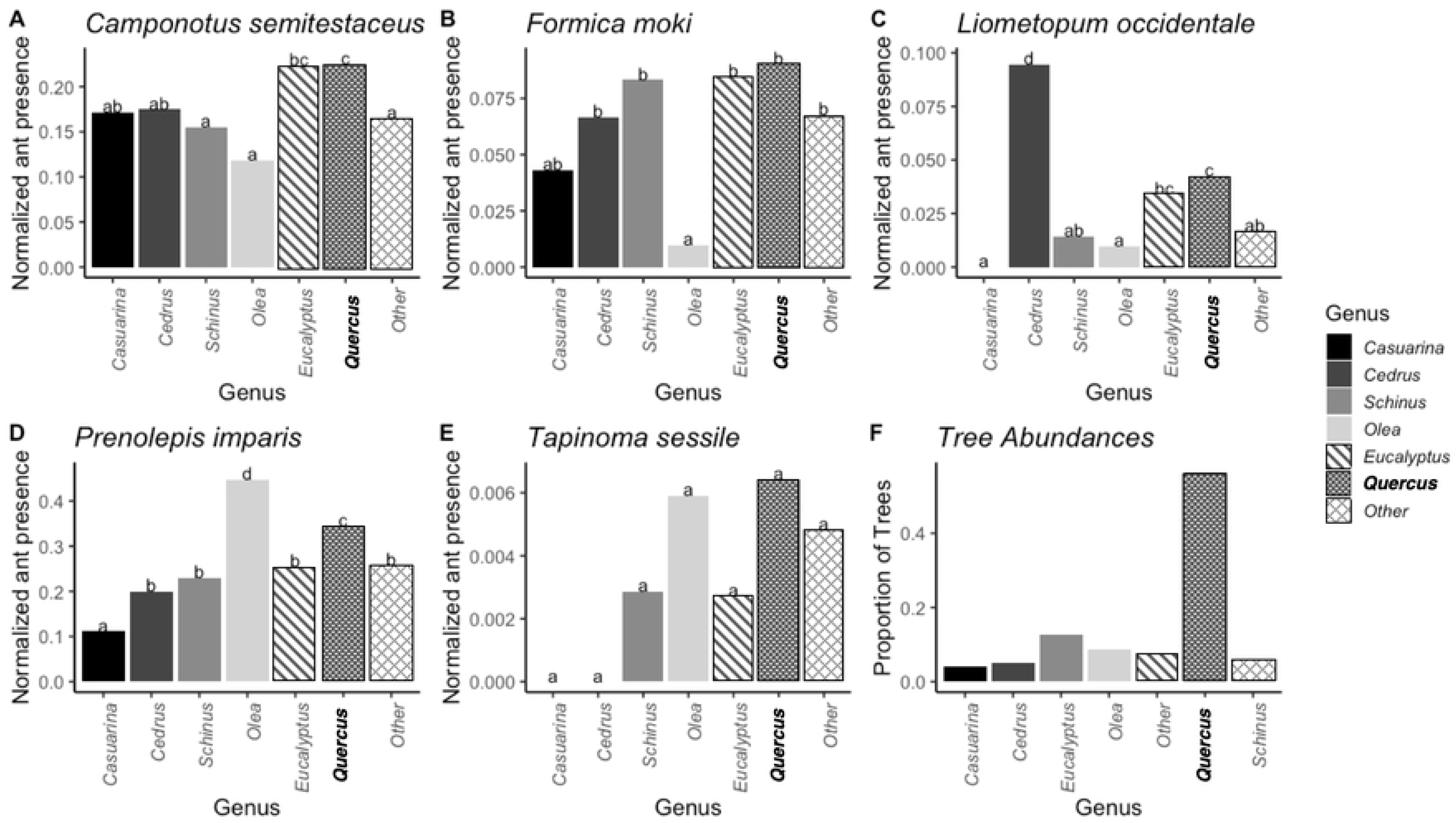
Ant presence by tree genus. Normalized ant presence is the number of trees on which an ant species was present divided by the total number of trees surveyed. Letters show significant differences in Dunn tests at the p < 0.05 level. Only tree genera with at least 50 individuals present in the study area were included separately; all other genera were included as “Other.” Panel F shows the proportion of each tree genus present. *Quercus* (in bold) is the only native genus among the six most abundant tree genera at the site.

Ants were observed significantly more often in native than non-native trees (t(7) = 6.2, p = 0.0002, Figure 3A). Ants were also observed significantly more often in evergreen than deciduous trees (t(7) = −11.3, p = 4.0 x 10^-6^, Figure 3B).

**Figure 3.**
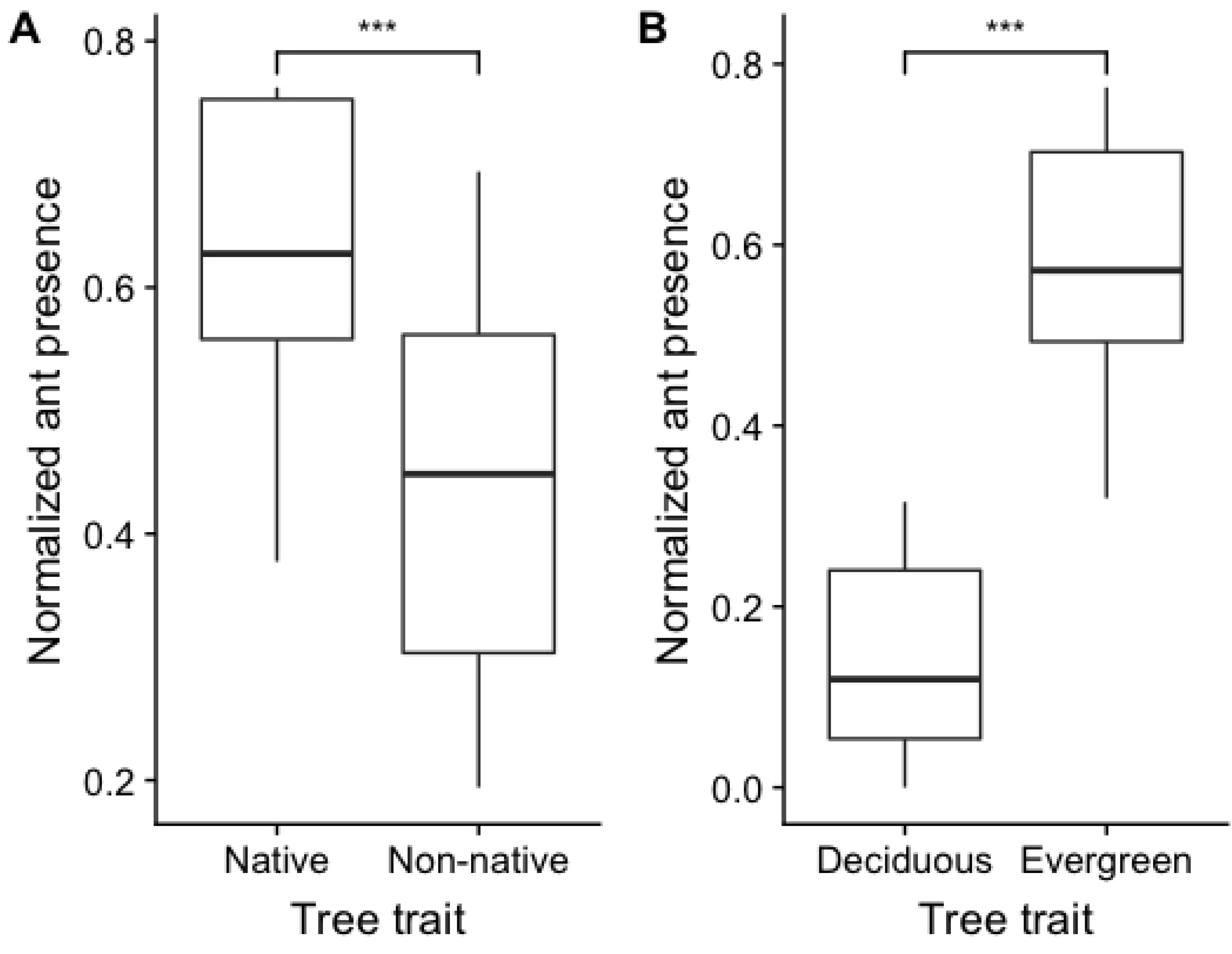
Normalized ant presence in native versus non-native trees (A) and deciduous versus evergreen trees (B). Normalized ant presence is the number of deciduous, evergreen, native, or non-native trees on which any ant species was present divided by the total number of that type of tree surveyed. *** indicates significance at the p < 0.001 level.

## Discussion

Our study suggests that niche partitioning is occurring in a mixed oak-exotic woodland ant community along three different axes: season, day/night, and tree genus. While other studies have investigated seasonal partitioning or day/night partitioning separately, our study demonstrates that seasonal and day/night scales may interact to influence partitioning.

**Figure 4.**
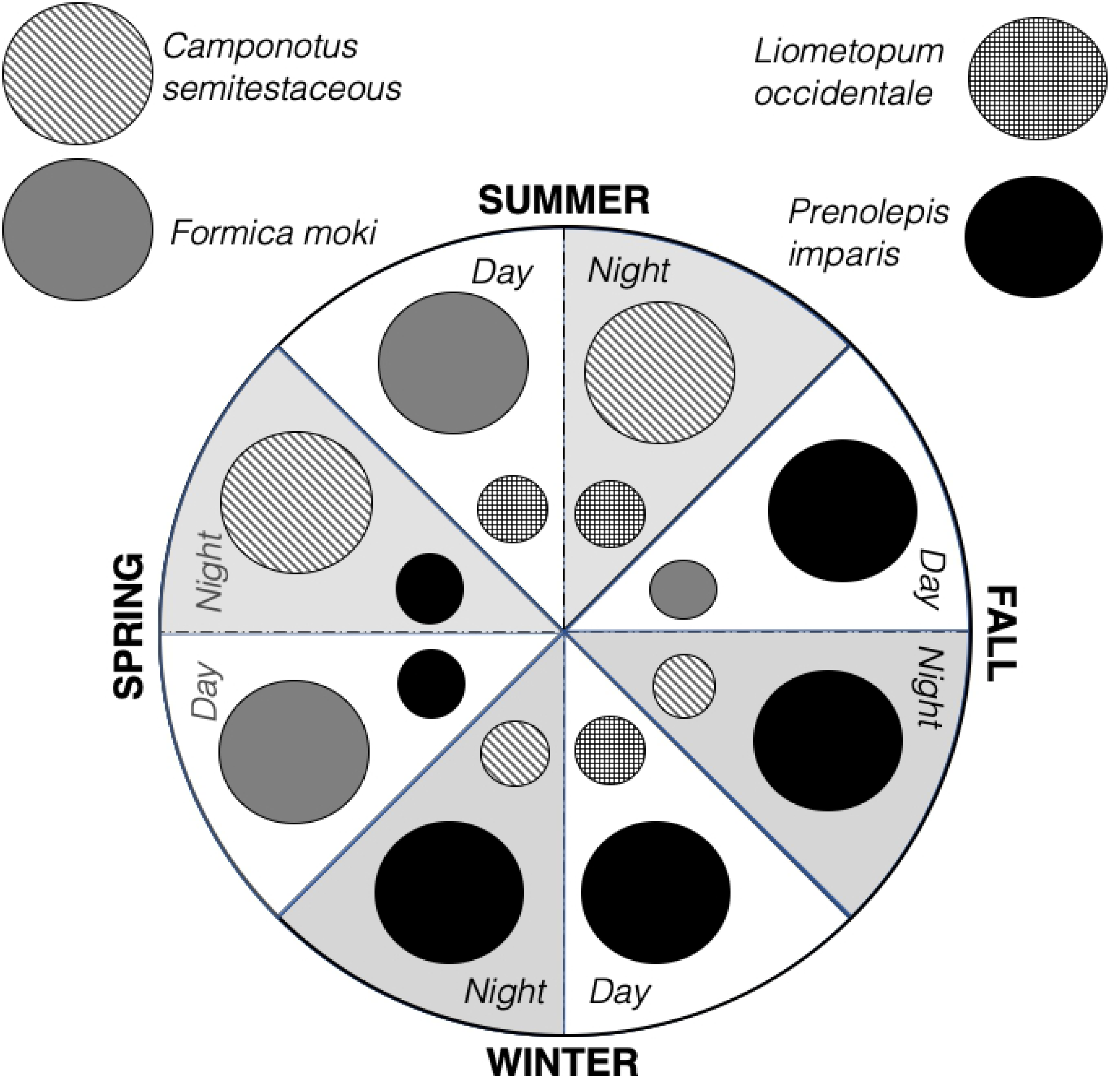
Differences in diel and seasonal ant activity among the most abundant ant species in a woodland ant community suggest niche partitioning. *Camponotus semitestaceus* was most active during spring and summer nights; *Formica moki* was most active during spring and summer days, and *Prenolepis imparis* was most active in fall and winter during both day and night. *Liometopum occidentale* was the second most active species during summer day, summer night, and winter day.

Thermal niche partitioning may account for the seasonal and diel patterns we observed. *Prenolepis imparis* foraged all day during fall, winter, and spring, which suggests this species relies on colder temperatures. Other studies have shown temperature-dependent activity in *P. imparis*; in other locations, *P. imparis* forages nocturnally in warm months and diurnally in cold months (10,30). *Formica moki* is active only during the day in warmer months, which could indicate a preference for warmer temperatures. By contrast, *Camponotus semitestaceus* has been observed to be nocturnal regardless of temperature or season (24). As in previous studies (10,31,32), we did not observe strong diel or seasonal activity patterns in *Liometopum occidentale* and *Tapinoma sessile*.

All species we observed foraged more frequently on native trees, particularly *Quercus* species. Some niche partitioning, however, occurred in exotic tree genera: *P. imparis* foraged frequently on *Olea*, and *L. occidentale* foraged most frequently on *Cedrus*, while other species avoided these two exotic tree genera. All species preferred to forage on evergreen trees independently of their preference for native trees, suggesting the distribution of honeydew resources may differ between evergreen and deciduous trees. *Quercus agrifolia*, a native, evergreen species, was the most common tree at the field site and may play an important role in maintaining ant biodiversity.

Species differed in trail formation and in their spatial use of trees (Figure S1), which may facilitate niche partitioning. *C. semitestaceus* formed diffuse trails somewhat frequently (about 30% of observations) and foraged from loosely clumped nest aggregations in the ground (Figure S1). *F. moki* rarely formed trails (Figure S1) and was usually present in low abundance (Figure S3).

*L. occidentale* frequently formed large foraging trails (~60% of observations, Figure S1), and foraged in clumped trail networks of around three to five trees, which often included a nest tree.

Several of the ant species, including *F. moki*, *L. occidentale*, and *P. imparis*, have been described as dominant based on bait assays (9,10,17,26,33). Though we did not offer baits, we frequently observed these species foraging together, which does not support a dominance-discovery tradeoff (34). The ants we observed were probably foraging on trees for aphid honeydew, a spatially stable food source for which repeated discovery may not be needed. Parr & Gibb (2012, 35) suggest that ant communities that rely on honeydew do not demonstrate dominance-discovery tradeoffs.

Further research is needed to explore the degree of dietary overlap for these species and whether the limitation and distribution of honeydew resources influences ant community structure. Preferences for particular tree genera may be driven by differences among tree genera in aphid species composition and honeydew availability. The emerging threat of sudden oak death in *Quercus* may affect ant communities because these species may be disproportionately important in supporting honeydew-feeding ants. Thermal niche partitioning may have important consequences for coexistence after climate change.

## Acknowledgements

Authors DM, RN and DMG contributed equally to the writing of this manuscript. This project was financially supported by a Small Grant awarded by the Stanford University Office of Undergraduate Advising and Research. We gratefully acknowledge Angela Gu, Duncan Coleman, Eleanor Glockner, Julie Fukunaga, Katie Lan, and Miranda Vogt for many hours of field work assistance. We thank Rodolfo Dirzo, Daniel Friedman and Maria Wojakowski for providing advice and assistance with statistical analysis. Leander D.L. Anderegg, Talia Borofsky, Kaleda Denton, and Tyler McFadden provided helpful comments on the manuscript. The Stanford Maps and Records Office (340 Bonair Siding, Stanford, CA 94305) provided detailed maps of trees on the Stanford grounds that were invaluable for this project.

